# Immune escape and replicative capacity of Omicron lineages BA.1, BA.2, BA.5.1, BQ.1, XBB.1.5, EG.5.1 and JN.1.1

**DOI:** 10.1101/2024.02.14.579654

**Authors:** Meriem Bekliz, Manel Essaidi-Laziosi, Kenneth Adea, Krisztina Hosszu-Fellous, Catia Alvarez, Mathilde Bellon, Pascale Sattonnet-Roche, Olha Puhach, Damien Dbeissi, Maria Eugenia Zaballa, Silvia Stringhini, Idris Guessous, Pauline Vetter, Christiane S Eberhardt, Laurent Kaiser, Isabella Eckerle

**Affiliations:** Department of Medicine, University of Geneva, Geneva, Switzerland.; Geneva Centre for Emerging Viral Diseases, University Hospitals of Geneva and University of Geneva, Geneva, Switzerland; Division of Infectious Diseases, Geneva University Hospitals, Geneva, Switzerland; Unit of Population Epidemiology, Division of Primary Care Medicine, Geneva University Hospitals, Geneva, Switzerland; Department of Health and Community Medicine, Faculty of Medicine, University of Geneva, Geneva, Switzerland; University Center for General Medicine and Public Health, University of Lausanne, Lausanne, Switzerland; Center for Vaccinology and Neonatal Immunology, Department of Pathology and Immunology, University of Geneva, Geneva, Switzerland; Division of General Pediatrics, Department of Woman, Child and Adolescent Medicine, Faculty of Medicine, University of Geneva, Geneva, Switzerland; Center of Vaccinology, University Hospitals of Geneva, Geneva, Switzerland

**Keywords:** SARS-CoV-2, omicron variants, viral replication, immune escape, neutralisation

## Abstract

In the 5^th^ year after the emergence of SARS-CoV-2, Omicron lineages continue to evolve and cause infections. Here, we used eight authentic SARS-CoV-2 isolates to assess their capacity to escape immunity of different exposure histories and their replicative capacity in polarized human airway epithelial cells (HAE) derived from the nasal and bronchial epithelium.

Using live-virus neutralization assays of 108 human sera or plasma of different immunological backgrounds, progressive immune escape was observed from B.1 (ancestral virus) to EG.5.1, but no significant difference between EG.5.1 and JN.1.1. Vaccinated individuals without natural infection and individuals with a single infection, but no vaccination showed markedly reduced or completely lost neutralization against the latest variants, while in those with hybrid immunity almost all sera showed some neutralization capacity. Furthermore, although absolute titers differed between groups, the pattern of immune escape between the variants remains comparable with strongest loss of neutralization observed for the latest variants.

*In vitro* studies with HAE at 33°C and 37°C showed some, but minor differences in virus replication and innate immune responses upon infection. Notably, infection with XBB.1.5, EG.5.1 and JN.1.1 showed slightly increased viral growth in nasal HAE at 33°C.

Altogether, these data underscore increasing immune escape across heterogeneous immunological backgrounds with gradually increasing antibody escape of evolving Omicron lineages until variant EG.5.1, but not any further for the latest dominant lineage JN.1.1. They also suggest that viral dynamics within Omicron lineages are driven by a combination of immune evasion and increase in viral replication.

## Introduction

The emergence of the SARS-CoV-2 in late 2019 has been followed by emergence of numerous virus variants [1]. Two years later, the emergence of the Omicron variant of concern (VOC) marked a significant shift, enhancing immune evasion due to extensive mutations in its Spike region. This has led to widespread infections by overcoming immunity from vaccines and prior natural infection. Omicron’s distinct genetic profile, particularly in the Spike region, positions it as a potential new serotype of SARS-CoV-2 [2, 3].

Global observations have documented successive waves of infections since the initial emergence of Omicron, with a huge number of evolving Omicron lineages. Consequently, a complex immunological landscape now exists in the population, where immunity is derived from vaccines, one or more infections due to pre-Omicron or Omicron variants, or a combination therefore, known as hybrid immunity. In most individuals in high income countries, the immunological background of SARS-CoV-2 today consists of multiple vaccinations and multiple exposures to SARS-CoV-2 through natural infection. On the other hand, some individuals have never received a vaccine and were only infected after the circulation of Omicron, in particular younger children, for which there is no vaccine recommendation in many countries. This leads to a heterogeneous immunological background in the population, ranging from both extremes of the exposure spectrum [2–9]. Despite background immunity in the population, SARS-CoV-2 continues to circulate. Multiple new variants have arisen within the Omicron clade, with XBB.1.5, XBB.1.16, EG.5, BA.2.86 and JN.1, being the latest variants of interest (VOIs) designated by WHO. BA.2.86, first identified in August 2023, has a remarkable number of mutations, known to allow antibody evasion, compared to earlier Omicron variants [10]. BA.2.86 did not show strong epidemiological signs of spread, but its decedent did, after BA.2.86 acquired an additional mutation S:L455S in the Spike, which became JN.1. This mutation was shown to be associated with significantly enhanced immune evasion capabilities but also increased transmissibility [5, 11–15]. In early 2024, JN.1 and its descendant lineages showed a strong increase globally, outcompeting earlier variants, and remains dominant as of mid 2024.

With reduced testing and surveillance and a highly variable immunological background in the population, obtaining data from epidemiology and/or clinical specimen collections has become more challenging. Therefore, an experimental assessment of immune escape and replicative capacity of emerging Omicron lineages *in vitro* is interesting, although it does not fully reflect the *in vivo* situation. Here, we investigated these aspects in SARS-CoV-2 Omicron lineages: BA.1, BA.2, BA.5.1, BQ.1, XBB.1.5, EG.5.1 and JN.1.1 compared to the ancestral SARS-CoV-2 B.1, using live viruses as a widely acknowledged gold standard, first, to compare immune escape capacities using sera from individuals with different immunological backgrounds by neutralization assays and second, to study the infection in relevant primary polarized airway epithelial culture models of the upper and lower respiratory tract.

## Methods

### Viruses and cells

Vero-E6 (ATCC CRL-1586) and Vero-E6-TMPRSS (Vero-E6 overexpressing TMPRSS2 protease, provided by National Institute for Biological Standards and Controls, NIBSC, Cat. Nr. 100978) cells were cultured as previously described [2, 16, 17]. All SARS-CoV-2 viruses used in this study were isolated from anonymized nasopharyngeal swabs collected at University Hospitals of Geneva (HUG) under an approval that allows the usage of anonymized left-over materials for virus culture. For this study, the following virus isolates were used (according to Pango lineages designation [18]): B.1 (ancestral SARS-CoV-2) and BQ.1, isolated and propagated on Vero-E6; BA.1 and BA.5.1, isolated on Vero-TMPRSS, then propagated to Vero-E6; BA.2, XBB.1.5, EG.5.1 and JN.1.1, isolated and propagated on Vero-TMPRSS. Both the initial clinical specimen and the obtained virus isolates were fully sequenced (**Table S1**). All virus stocks were titrated on the same cell line on which the virus stock was produced (either Vero-E6 or Vero-TMPRSS).

### SARS-CoV-2 infections in HAE

Infections with SARS-CoV-2 Omicron lineages BA.1, BA.2, BA.5.1, BQ.1, XBB.1.5, EG.5.1 and JN.1.1 were performed at 37°C or 33°C at 5% CO_2_ at a multiplicity of infection of approximately 0.1 in commercially available polarized tissues “MucilAir^TM^” (Epithelix SARL), *in vitro* reconstituted from human nasal or bronchial (3 donors from each group) epithelial cells of adult healthy donors cultured in an air-liquid interface (ALI) system, as previously described [16, 19, 20]. Viral replication was assessed at 24, 48, 72 and 96 hours post infection (hpi), as previously described [16, 19, 20].

### Assessment of host gene response

Induction of interferons IFN-α and IFN-β IFN-λ, ISG15 (Interferon stimulated gene 15), angiotensin-converting enzyme 2 (ACE-2) and Transmembrane Serine Protease 2 (TMPRSS2) was assessed by semi-quantitative real-time PCR for intracellular RNA collected at 96 hpi, as previously described [16, 19, 20].

### Human serum and plasma samples

Immunocompetent and healthy individual samples consisted of serum or plasma samples collected after vaccination, infection or a combination of both (hybrid immunity). Plasma or serum samples from vaccinated healthy individuals, vaccinated either with two or three doses (boosted) of BNT162b2 (Pfizer/BioNTech) or mRNA-1273 (Moderna) were available from a prospective observational studies (Ethics approval number: CCER 2021-00430 and CCER 2020-02323). Asymptomatic or undetected infections of the vaccinated-only group were excluded in those samples by testing all specimens for antibodies against SARS-CoV-2 nucleocapsid (Roche Elecsys anti-SARS-CoV-2 N assay). Specimens of individuals with hybrid immunity were collected from adults vaccinated with BNT162b2 or mRNA-1273 vaccines and with one or more documented infections. Breakthrough infection samples were collected from individuals with either 2x or 3x mRNA vaccination, followed by an Omicron BA.1 or BA.2 breakthrough infection, respectively (Ethics approval number: CCER 2020-02323). Serum samples from individuals with XBB breakthrough infection had received either 2x (n=2), 3x (n=7) or 4x (n=2) mRNA vaccination, followed by breakthrough infection with one of the XBB sublineages that occurred between March-June 2023 (specimens were left-over samples from a prospective observational study, ethics approval number: CCER 2022-01722). Four of these individuals had a subsequent infection prior to XBB infection. One of the individuals vaccinated with 4x mRNA had received a bivalent vaccine. For individuals with infections from two different VOCs (e.g. Alpha and Omicron or Delta and Omicron) had received either 1x (n=6), 2x (n=10) or 3x (n=2) mRNA vaccine (Ethics approval number: CCER 2020-02323). Convalescent sera from unvaccinated adults and children with confirmed SARS-CoV-2 infection in early 2022 were also available (Ethics approval number: CCER 2020-00881). Based on the data from our Swiss national genomic surveillance, the vast majority of the variants circulating at that time was Omicron BA.1 and BA.2, with only few Delta sequences remaining in January 2022 [21]. The infecting variant of each episode was either determined by sequencing of the diagnostic samples or extrapolated by the time of infection according to the information that was self-reported by the participant and/or by their parent (for children), taking the knowledge on variant circulation generated by the Swiss national genomic surveillance program into account [22]. Data on collection time of specimens after vaccination and/or infection are displayed in **Tables 1-5**. Only one serum per individual of a single collection time point was used in this study.

**Table 1.** Characteristic of vaccinated individuals’ samples.

| **Vaccine** | **Number of individuals** | **Sample type** | **Gender**<br>**(M/F)** | **Age**<br>**mean** | **WPLV**<br>**Mean weeks (range)** | **Date of last vaccination** |
| --- | --- | --- | --- | --- | --- | --- |
| **2x mRNA vaccine** | 14 | Plasma | 2/12 | 51 (36-62) | 4 (4-5) | March-May 2021 |
| **3x mRNA vaccine** | 19 | Serum | 9/10 | 43 (26-63) | 13 (3-22) | November 2021-January 2022 |

A written informed consent was obtained from all adult participants, and from the legally appointed representatives (parents) of all minor participants. All necessary approvals were obtained from the Cantonal Ethical board of the Canton of Geneva, Switzerland (Commission Cantonal d’Ethique de la Recherche, CCER). Since no differences are to be expected in neutralizing capacity between plasma or serum, both sample types were used in parallel.

### Focus reduction neutralization test (FRNT)

FRNT was used to determine the infectious titer after neutralization. Vero-TMPRSS cells were seeded at a density of 4 × 10^5^ cells/mL in 96-well cell culture plates. All sera/plasma and Vero-TMPRSS cells infections were prepared as previously described [2]. After incubation for 16-24 hours at 37 °C, 5% CO_2_, the plates were fixed and stained for SARS-CoV-2 nucleocapsid protein as described previously [17, 23]. The 90% reduction endpoint titers (FRNT_90_) were calculated as previously described [2]. For samples that did not reach 90% reduction at a 1:10 dilution, we extrapolated the titer until a dilution of 0.5. If the extrapolation reached a titer below 0.5, the sample was given a value of 0.5. All samples with a titer below 1, i.e. undiluted sample are considered negative.

Data was recorded in Excel 2019. Geometric means with 95% CI were used for the comparison of FRNT_90_ titers. Statistical analyses for FRNT_90_ were conducted using GraphPad Prism version 9.1.0 software and performed using repeated measures one-way ANOVA with Dunnett’s multiple comparisons test with log_10_ transformed FRNT_90_ titers.

## Results

### 1. Virus neutralization to Omicron lineages of sera or plasma after infection, vaccination, and hybrid immunity

A panel of sera/plasma was used from: (i) double or monovalent boosted vaccinated individuals with BNT162b2 or mRNA-1273 without prior or subsequent infection (**Table 1**); (ii) individuals with hybrid immunity after receiving two or three doses of mRNA vaccine followed by a break-through infection with either Omicron BA.1 or BA.2, respectively (**Table 2**); (iii) individuals with hybrid immunity due to XBB-variant breakthrough infection (**Table 3**); (iv) individuals with hybrid immunity receiving between one to three doses of mRNA vaccine and two different documented VOCs infections (either infection with Alpha or Delta, followed by infection with Omicron) (**Table 4**); and (v) a panel of convalescent sera from unvaccinated individuals, adults and children infected between January and March 2022, a period that was characterized by high circulation of Omicron BA.1 and BA.2 (**Table 5**).

**Table 2.** Characteristic of hybrid immunity (breakthrough with BA.1 or BA.2) individuals’ samples.

| **Vaccination/infection status** | **Number of individuals** | **Sample type** | **Gender**<br>**(M/F)** | **Age**<br>**mean** | **WPLV**<br>**Mean weeks (range)** | **WPLP**<br>**Mean weeks (range)** | **Interval vaccination – infection**<br>**Mean weeks (range)** | **Variant**<br>**Identification<sup>1</sup>** |
| --- | --- | --- | --- | --- | --- | --- | --- | --- |
| **2x mRNA vaccine & BA.1 breakthrough** | 11 | Serum | 5/6 | 39 (25-55) | 34 (12-52) | 15 (2-15) | 27 (8-41) | Sequencing (n=11) |
| **3x mRNA vaccine & BA.2 breakthrough** | 11 | Serum | 5/6 | 39 (25-62) | 22 (15-35) | 7 (3-10) | 15 (5-32) | Sequencing (n=11) |

**Table 3.** Characteristic of hybrid immunity (breakthrough with XBB) individuals’ samples.

| **Vaccination<sup>2</sup>/infection status<sup>3</sup>** | **Number of individuals** | **Sample type** | **Gender (M/F)** | **Age mean** | **Number of vaccinations** | **WPLV Mean weeks (range)** | **WPLP Mean weeks (range)** | **Variant<sup>1</sup> identification** |
| --- | --- | --- | --- | --- | --- | --- | --- | --- |
| **Vaccination & XBB breakthrough infection** | 11 | Serum | 1/10 | 44 (24-61) | 2, 3 or 4 | 73 (36-93) | 10 (2-19) | Sequencing (n=11) |

**Table 4.** Characteristic of multi-VOC infected individuals’ samples.

| **Vaccination<sup>4</sup>/infection status** | **Number of individuals** | **Sample type** | **Gender (M/F)** | **Age mean** | **WPLV Mean weeks (range)** | **WPLD Mean weeks (range)** | **Interval vaccination – last infection Mean weeks (range)** | **Variant identification<sup>1</sup>** |  |
| --- | --- | --- | --- | --- | --- | --- | --- | --- | --- |
|  |  |  |  |  |  |  |  | **Alpha or Delta** | **Omicron** |
| **Vaccination + Alpha & Omicron breakthrough** | 9 | Serum | 5/4 | 41 (25-50) | 42 (30-64) | 17 (10-29) | 25 (12-44) | Sequencing (n=3) | Sequencing (n=5) |
| **Vaccination + Delta & Omicron breakthrough** | 9 | Serum | 3/6 | 40 (29-51) | 53 (27-87) | 21 (4-41) | 31 (18-55) | Sequencing (n=6) | Sequencing (n=1) |

**Table 5.** Characteristic of convalescent children and adults’ individuals’ samples.

| **Infecting virus<sup>5</sup>** | **Number of individuals** | **Sample type** | **Gender**<br>**(M/F)** | **Age**<br>**mean** | **WPLD**<br>**Mean days**<br>**(range)** | **Infection**<br>**period** | **Variant**<br>**Identification<sup>1</sup>** |
| --- | --- | --- | --- | --- | --- | --- | --- |
| **Omicron in adults** | 12 | Serum | 2/10 | 40 (28-46) | 15 (9-21) | January-March<br>2022 | Extrapolated<br>(n=12) |
| **Omicron in children** | 12 | Serum | 6/6 | 5 (2-7) | 17 (12-21) | January-March<br>2022 | Extrapolated<br>(n=12) |

*WPLV* weeks post last dose of vaccine; *WPLP* weeks post last positive RT-PCR; *WPLD* weeks post last positive diagnosis;
<sup>1</sup>The infecting variant of each episode was either determined by sequencing of the diagnostic samples or extrapolated according to the dominance of the variant at that time based on the data from our Swiss national genomic surveillance (**Figure S1**).
<sup>2</sup>Vaccination consisted of either 2 (n=2), 3 (n=7) or 4 (n=2) doses of BNT162b2 or mRNA-1273 vaccine. One of the individuals vaccinated with 4 doses of mRNA had received a bivalent vaccine.
<sup>3</sup>Infection consisted of either 1 (n=7) or 2 (n=4) infection including one of the Omicron XBB variants infection.
<sup>4</sup>Vaccination consisted of either 1 (n=6), 2 (n=10) or 3 (n=2) doses of BNT162b2 or mRNA-1273 vaccine.
<sup>5</sup>Individuals were infected between January and March 2022 when the predominant circulation of Omicron BA.1 and BA.2 were taking place (<90%).

#### 2.1. Neutralizing capacity in vaccinated but never-infected individuals against Omicron lineages

We investigated a total of 33 individuals’ specimens, either double-vaccinated (n=14) or monovalent boosted individuals that had never been infected with SARS-CoV-2 (n=19) for neutralization against ancestral SARS-CoV-2 lineage B.1 as well as SARS-CoV-2 Omicron lineages BA.1, BA.2, BA.5.1, BQ.1, XBB.1.5, EG.5.1 and JN.1.1. For double-vaccinated individuals, the highest neutralizing capacity was observed against the ancestral SARS-CoV-2 virus B.1 with geometric mean FRNT_90_ titers of 251.7 (95%CI: 158.3-400.1) but very reduced titers were observed against all Omicron lineages with FRNT_90_ titers of 5.7 (95%CI: 2.8-11.5) for BA.1, 10.4 (95%CI: 6.6-16.3) for BA.2, 2.9 (95%CI: 1.5-5.5) for BA.5.1, 1.5 (95%CI: 0.8-3.0) for BQ.1, 1.0 (95%CI: 0.6-1.4) for XBB.1.5 and 0.6 (95%CI: 0.5-0.8) for JN.1.1. None of the samples neutralized EG.5.1. Although titers were very reduced compared to B.1, none of the samples failed to neutralize Omicron BA.2. An increasing number of samples with complete failure to neutralize was observed for Omicron BA.1, BA.5.1, BQ.1, XBB.1.5, EG.5.1 and JN.1.1 with 2/14, 4/14, 7/14, 7/14, 14/14 and 12/14, respectively (**Figure 1A**).

**Figure 1.**
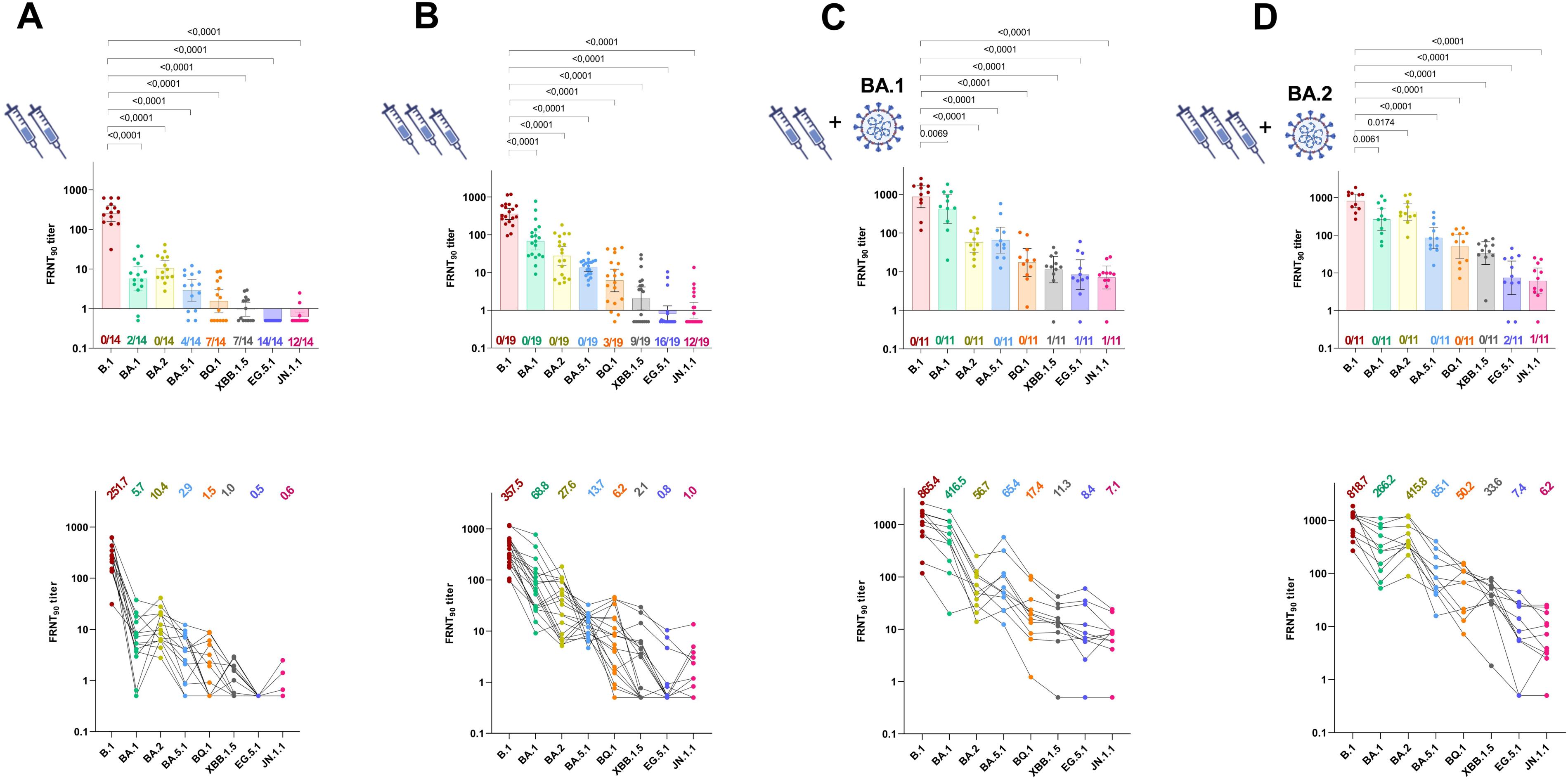
Neutralization in vaccine and hybrid immunity-derived blood specimens against eight authentic isolates of SARS-CoV-2 variants (B.1 and Omicron lineages including BA.1, BA.2, BA.5.1, BQ.1, XBB.1.5, EG.5.1 and JN.1.1). Bars represent geometric mean titers (GMT) of 90% reduction endpoint titers (FRNT_90_) with 95% confidence interval. **A**–**D** Cohorts of individuals with **A)** double-dose mRNA vaccination (n=14), **B)** boosted mRNA vaccination (n=19), **C)** BA.1 breakthrough infection following double mRNA vaccination (n=11) and **D)** BA.2 breakthrough infection following 3 mRNA vaccination (n=11). Coloured numbers below each bar represent the number of specimens with complete loss of neutralization (FRNT_90_ titer < 1). Repeated measures one-way ANOVA with Dunnett’s multiple comparisons test using log_10_ transformed FRNT_90_ titers was performed to analyze the statistical significance.

For boosted individuals, overall geometric mean FRNT_90_ titers were higher against all viruses. Geometric mean FRNT_90_ titers were 357.5 (95%CI: 255.3-500.6), 68.8 (95%CI: 39.7-119.2), 27.6 (95%CI: 15.5-49.1), 13.7 (95%CI: 10.7-17.4), 6.2 (95%CI: 3.1-12.3), 2.1 (95%CI: 1.0-4.1), 1.5, 0.8 (95%CI: 0.5-1.3) and 1.0 (95%CI: 0.6-1.6) against B.1, BA.1, BA.2, BA.5.1, BQ.1, XBB, EG.5.1 and, JN.1.1, respectively. No complete loss of neutralization in this group was observed for variants B.1, BA.1, BA.2, and BA.5.1, while 3/19 samples were not neutralized for BQ.1, 9/19 for XBB.1.5, 16/19 for EG.5.1 and 12/19 for JN.1.1 (**Figure 1B**).

#### 2.2. Neutralizing capacity of vaccinated individuals with breakthrough infection (hybrid immunity)

We investigated the impact of BA.1 breakthrough infection in individuals vaccinated with two doses of mRNA vaccine. Geometric mean FRNT_90_ titers in the hybrid immunity group were higher than for vaccinated individuals. They were 865.4 (95%CI: 450.9-1661.0) against B.1, 416.5 (95%CI: 175.6-987.8) against BA.1, 56.7 (95%CI: 31.8-100.9) against BA.2, 65.4 (95%CI: 30.3-141.4) against BA.5.1, 17.4 (95%CI: 7.6-39.9) against BQ.1, 11.3 (95%CI: 5.1-24.9) against XBB.1.5, 8.4 (95%CI: 3.5-20.3) against EG.5.1 and 7.1 (95%CI: 3.6-14.0) against JN.1.1. Complete loss of neutralization was observed only for 1/11 sample each for XBB.1.5, EG.5.1 and JN.1.1 (**Figure 1C**).

For boosted individuals with BA.2 breakthrough infection, geometric mean FRNT_90_ titers were 818.7 (95%CI: 541.5-1238.0) for B.1, 266.2 (95%CI: 134.2-528.0) for BA.1; 415.8 (95%CI: 250.5-690.1) for BA.2, 85.1 (95%CI: 44.7-161.9) for BA.5.1, 50.2 (95%CI: 24.2-104.0) for BQ.1, 33.6 (95%CI: 16.7-67.4) for XBB.1.5, 7.4 (95%CI: 2.7-20.7) for EG.5.1 and 6.2 (95%CI: 2.8-13.5) for JN.1.1. No complete loss of neutralization in this group was observed for variants B.1, BA.1, BA.2, BA.5.1, BQ.1 and XBB.1.5, while 2/11 samples were not neutralized for EG.5.1 and 1/11 for JN.1.1 (**Figure 1D**).

Although titers for B.1 were highest in both groups, the second highest neutralization titers were found against the infecting virus, *e.g.*: neutralization of BA.1 was higher in the BA.1-infected group and neutralization of BA.2 was higher in the BA.2.-infected group. No difference was seen between the groups in the neutralization titers for EG.5.1. and JN.1.1 which were both comparably low independent of the infecting virus.

We then investigated the impact of breakthrough infections in individuals (n=11) who have been infected with one of the Omicron XBB subvariants between March and June 2023. The geometric mean FRNT_90_ titers were 1984.0 (95%CI: 1109-3547) against B.1, 46.0 (95%CI: 18.2-116.4) against XBB.1.5, 31.8 (95%CI: 9.3-108.6) against EG.5.1 and 21.1 (95%CI: 9.1-48.7) against JN.1.1. All sera were able to neutralize in this group (**Figure 2A**).

**Figure 2.**
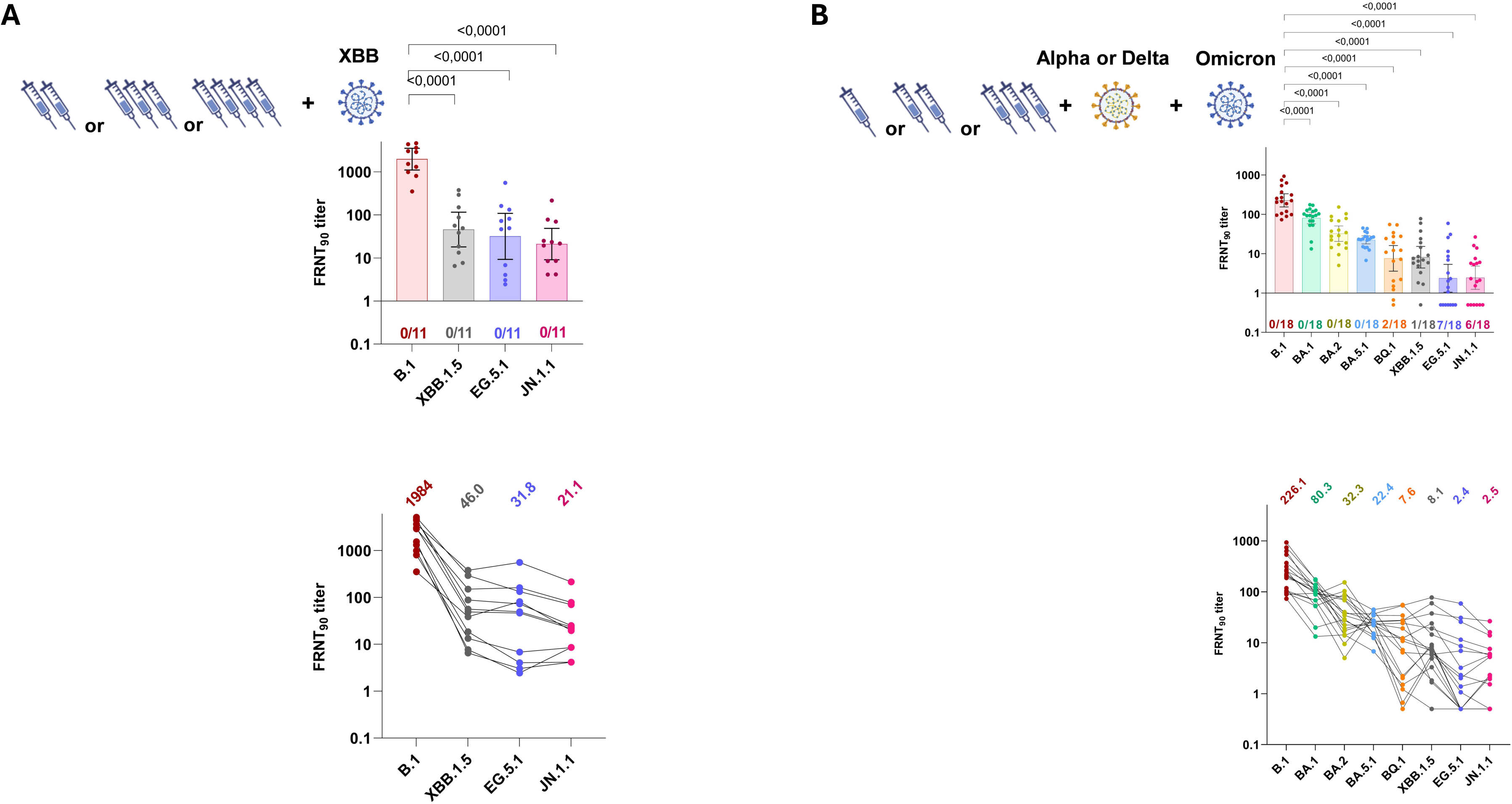
**A)** Neutralization in XBB exposure-derived blood specimens against four authentic isolates of SARS-CoV-2 variants (B.1 and Omicron lineages including XBB.1.5, EG.5.1 and JN.1.1). Cohort of specimens after XBB-derived infections with one of Omicron XBB variant. Bars represent geometric mean titers (GMT) of 90% reduction endpoint titers (FRNT_90_) with 95% confidence interval. Coloured numbers below each bar represent the number of specimens with complete loss of neutralization (FRNT_90_ titer < 1). Repeated measures one-way ANOVA with Dunnett’s multiple comparisons test using log_10_ transformed FRNT_90_ titers was performed to analyze the statistical significance. **B)** Neutralization in hybrid immunity-derived blood specimens against eight authentic isolates of SARS-CoV-2 variants (B.1 and Omicron lineages including BA.1, BA.2, BA.5.1, BQ.1, XBB.1.5, EG.5.1 and JN.1.1). Bars represent geometric mean titers (GMT) of 90% reduction endpoint titers (FRNT_90_) with 95% confidence interval. Cohort of vaccinated individuals with dual SARS-CoV-2 infections (e.g. Alpha and Omicron (n=9) or Delta and Omicron (n=9)). Coloured numbers below each bar represent the number of specimens with complete loss of neutralization (FRNT_90_ titer < 1). Repeated measures one-way ANOVA with Dunnett’s multiple comparisons test using log_10_ transformed FRNT_90_ titers was performed to analyze the statistical significance.

#### 2.3. Neutralizing capacity in vaccinated individuals with breakthrough infections from two antigenically different VOCs (multi-variant hybrid immunity) towards Omicron lineages

We investigated vaccinated individuals (n=18) who subsequently have been exposed to at least two antigenically different variants through two independent infection episodes that included a pre-Omicron VOC (Alpha, n=9 or Delta, n=9) and another infection episode with an Omicron lineage. The geometric mean FRNT_90_ titers were 226.2 (95%CI: 153.8-332.8) against B.1, 80.33 (95%CI: 57.3-112.6) against BA.1, 32.3 (95%CI: 20.5-50.7) against Omicron BA.2, 22.4 (95%CI: 17.7-28.4) against BA.5.1, 7.6 (95%CI: 3.6-16.1) against BQ.1, 8.1 (95%CI: 4.3-15.3) against XBB.1.5, 2.4 (95%CI: 1.0-5.4) against EG.5.1 and 2.5 (95%CI: 1.2-4.8) against JN.1.1. Loss of neutralization was observed for BQ.1, XBB.1.5, EG.5.1 and JN.1.1 in 2/18, 1/18, 7/18 and 6/18 samples, respectively (**Figure 2B**).

#### 2.4. Neutralizing capacity from unvaccinated adults and children infected in early 2022

We also studied neutralization of Omicron variants in 24 sera (adults, n=12; children, n=12) of unvaccinated individuals with a single infection between January and March 2022 (most likely exposed to BA.1 or BA.2). Due to the limited volume of serum available for this group, we only assessed neutralization towards Omicron variants BA.1, BQ.1, XBB.1.5, EG.5.1 and JN.1.1 in this cohort. Geometric mean FRNT_90_ titers in this group for adult individuals were 29.1 (95%CI: 13.0-65.1) against BA.1, 1.3 (95%CI: 0.6-2.6) against BQ.1; 0.8 (95%CI: 0.5-1.4) against XBB.1.5 and none of the samples neutralized EG.5.1 and Omicron JN.1.1. Complete loss of neutralization was observed for 7/12 samples for BQ.1, 8/12 samples for XBB.1.5 and all samples for EG.5.1 and JN.1.1 (**Figure 3A**). For children, geometric mean FRNT_90_ titers were 43.3 (95%CI: 18.0-103.7) against BA.1, 2.3 (95%CI: 0.9-5.8) against BQ.1, 1.1 (95%CI: 0.5-2.4) against XBB.1.5, 1.2 (95%CI: 0.6-2.5) against EG.5.1 and 0.6 (95%CI: 0.4-0.8) against JN.1.1. Of note, complete loss of neutralization was observed for 5/12 samples for BQ.1, 8/12 samples for XBB.1.5, 6/12 samples for EG.5.1 and 11/12 for JN.1.1 (**Figure 3B**).

**Figure 3.**
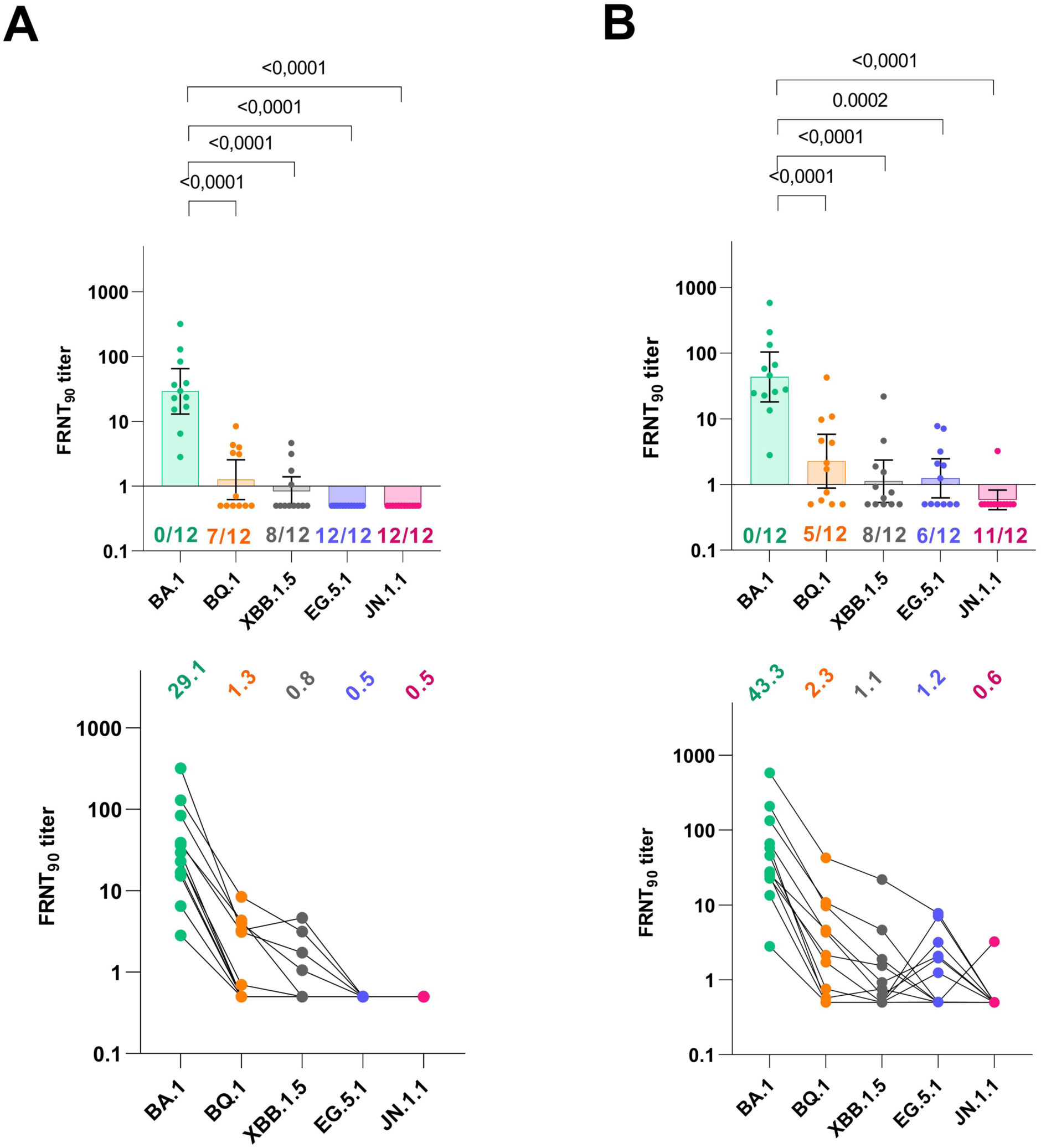
Neutralization in infection-derived blood specimens against five authentic isolates of SARS-CoV-2 variants (omicron lineages including BA.1, BQ.1, XBB.1.5, EG.5.1 and JN.1.1). Bars represent geometric mean titers (GMT) of 90% reduction endpoint titers (FRNT_90_) with 95% confidence interval. Cohorts of convalescent specimens that are derived from **A)** unvaccinated adult individuals (n=12) and **B)** unvaccinated children individuals (n=12) with confirmed SARS-CoV-2 infection in early 2022 (thus, probably Omicron BA.1 or BA.2). Coloured numbers below each bar represent the number of specimens with complete loss of neutralization (FRNT_90_ titer < 1). Repeated measures one-way ANOVA with Dunnett’s multiple comparisons test using log_10_ transformed FRNT_90_ titers was performed to analyze the statistical significance.

**Figure 4.**
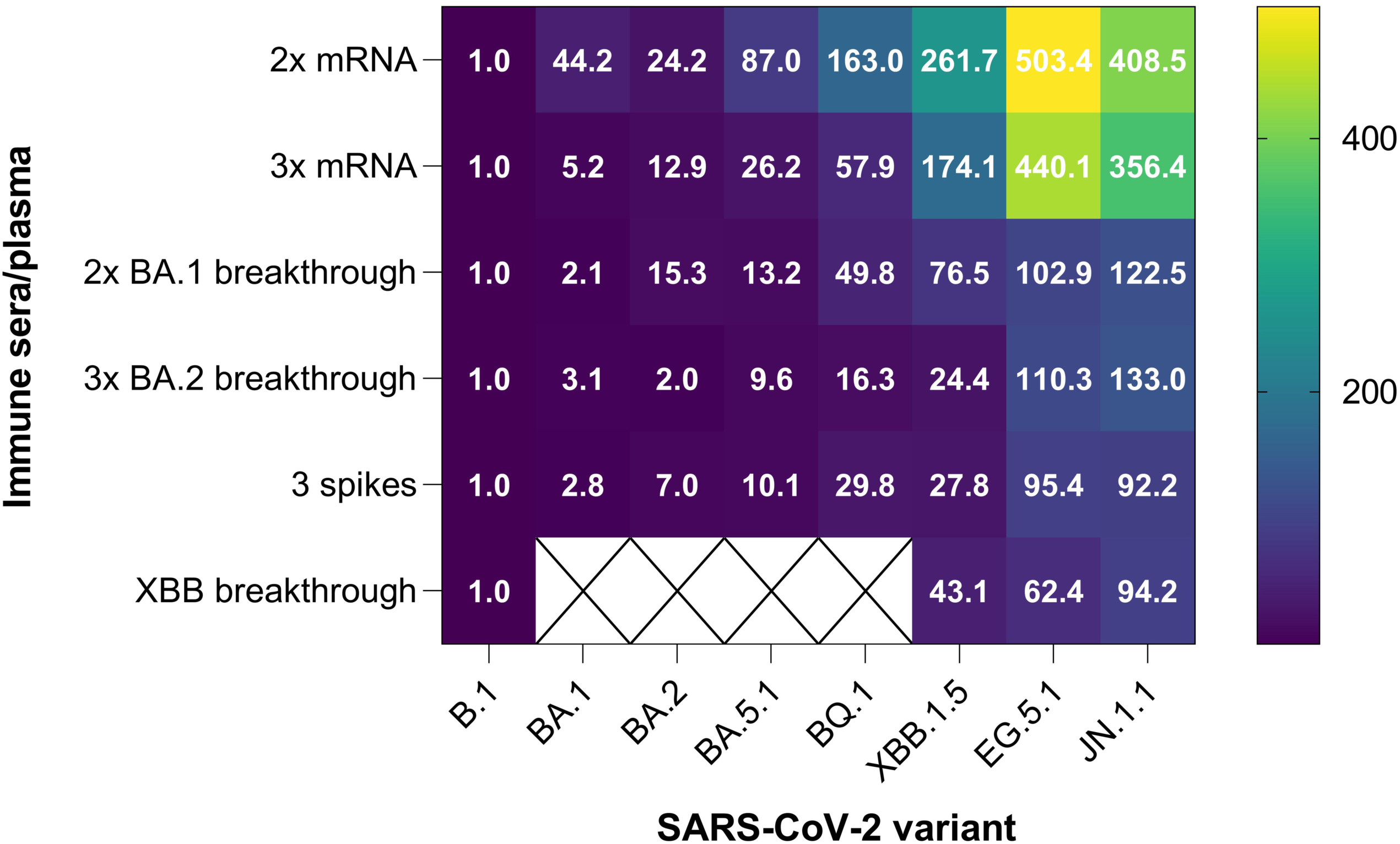
Heatmap of fold-reduction in neutralization based on FRNT_90_ data. Values of fold-reduction in neutralization (FRNT_90_) of B.1 and Omicron sublineages including BA.1, BA.2, BA.5.1, BQ.1, XBB.1.5, EG.5.1 and JN.1.1 are presented as heat maps with lighter colors implying greater changes. The immune sera/plasma were organized into cohorts of individuals with double-dose mRNA vaccination (n=14), boosted individuals with three doses of mRNA vaccine (n=19), BA.1 breakthrough infection of double-vaccinated individuals (n=11), BA.2 breakthrough infection individuals following 3 doses of mRNA vaccine (n=11), vaccinated individual with XBB breakthrough infection (n=11) and vaccinated Individuals with dual SARS-CoV-2 Infections (n=18).

**Figure 5.**
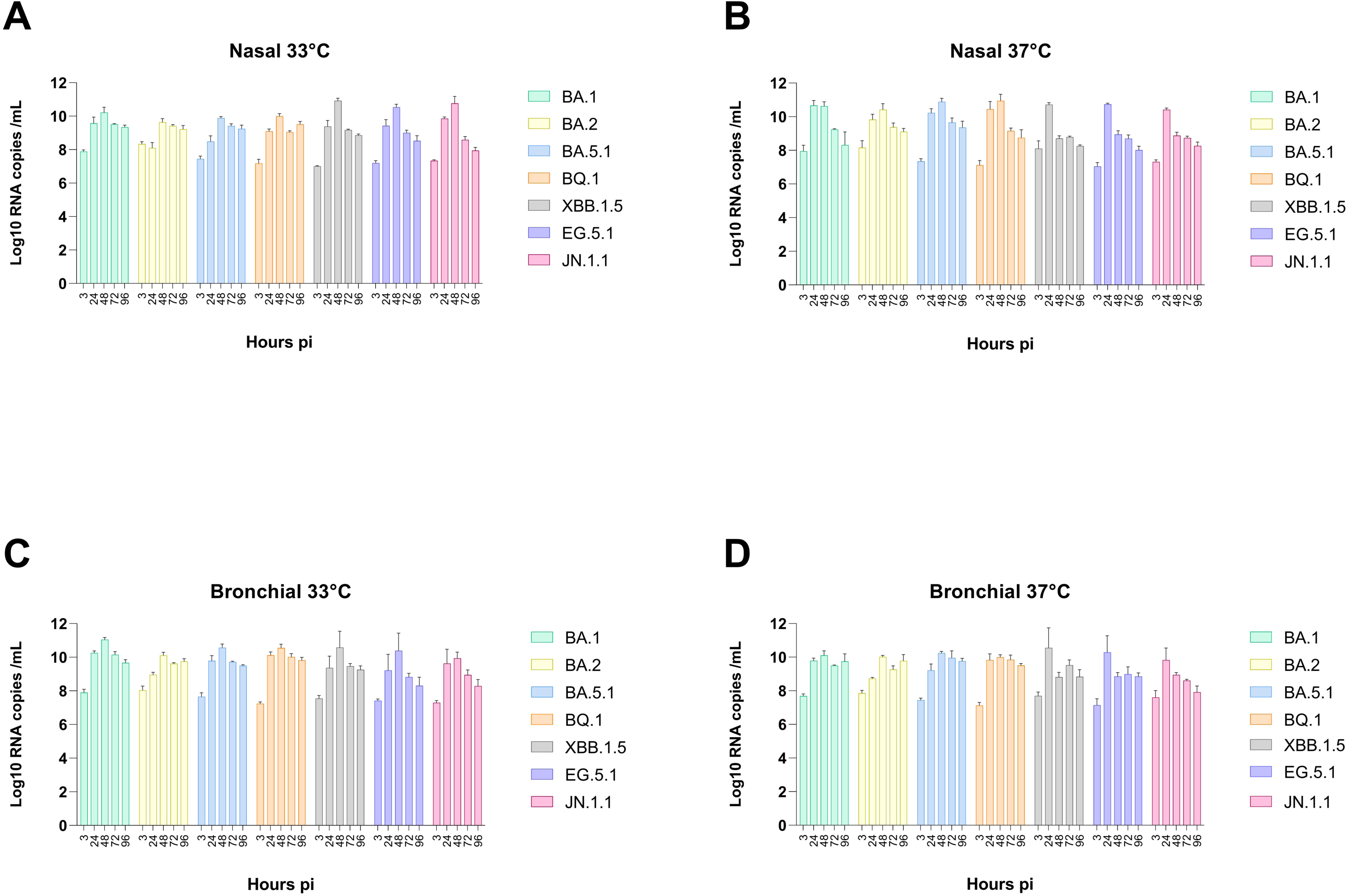
Replication of SARS-CoV-2 Omicron lineages in HAE. Nasal (**A** and **B)** and Bronchial (**C** and **D**) HAE were infected with SARS-CoV-2 Omicron lineages BA.1, BA.2, BA.5.1, BQ.1, XBB.1.5, EG.5.1 and JN.1.1 at 33°C (**A** and **C**) and 37°C (**B** and **D**). Viral replication was assessed by the quantification of viral RNA from apical washes collected at 3hpi (baseline), 24hpi, 48hpi, 72hpi and 96hpi. For each cell origin, HAE from 3 donors have been tested. Data are expressed as mean of the log of viral RNA copies/mL (log_10_ RNA/mL) and SEM.

#### 2.5. Heatmap of neutralization data across different immunological backgrounds

To summarize the findings across the cohorts, we have displayed the fold change of geometric mean FRNT_90_ titers in comparison to the ancestral virus B.1 for all cohorts (**Figure 3**). Across cohorts, a consecutive loss in neutralization was observed from B.1 to BA.1/BA.2. to BA.5.1 to BQ.1 to XBB.1.5. and to EG.5.1/JN.1. The effect was the strongest for sera from individuals that were only vaccinated but never infected, and the least pronounced for individuals exposed to more than one natural infection with different variants. The differences between EG.5.1 and JN.1. were only subtle across groups, and JN.1.1 did not show an enhanced immune escape compared to EG.5.1. In vaccinated but never infected individuals, there was even a tendency for better neutralization of JN.1.1 compared to EG.5.1.

### 2. Replicative capacity and innate immune responses of Omicron sublineages in nasal and bronchial HAE

To understand if Omicron lineages differ in their ability to replicate, we infected polarized HAE of nasal (3 donors) and bronchial (3 donors) origin at different temperatures (**Figure 1**) with different Omicron lineages. Comparison across lineages overall revealed similar kinetics and replication range, with a rapid increase in viral RNA, reaching peak viral loads at 24/48hpi followed by a slight decline at 72 and 96hpi. An increase in viral RNA was observed at 24h for almost all lineages under all conditions. In nasal HAE, virus replication was slightly lower in the physiological conditions (33°C) than at 37°C, where the peak of replication, reached at 48h, varied from 9.6 log10 RNAc/mL (for BA.2), to 10.9 log10RNA c/mL (for XBB). The most recent subvariants XBB.1.5, EG.5.1 and JN.1.1 showed better replication efficacy (higher than 10.5 log10 RNAc/mL). At 37°C, the peak was reached at 1dpi, except for BA.2, BA.5.1 and BQ.1. Similar patterns were found in bronchial HAE with comparable levels of replication at both temperatures. BA.2 again showed the lowest level of replication (10.1 log10 RNAc/mL at 33°C and 10.0 log10 RNAc/mL at 37°C). The three most recent subvariants all showed earlier replication peaks with a stronger increase of viral RNA at 37°C.

To compare innate immune responses after infection between Omicron lineages, we studied IFN-α and -β and IFN-λ interferon responses and the induction of downstream interferon-stimulated genes 15 (ISG15) at the end of the infection experiments (96hpi) in the infected and non-infected HAE cultures (**Figure 6**). IFN-α by all variant’s induction was barely observed (less than 1log increase versus non-infected cells) at 33°C in infected nasal and bronchial HAE and at 37°C in only in bronchial tissues. For all IFNs and ISG15, the lowest and highest inductions were found in infected nasal HAE at 33°C and bronchial HAE at 37°C, respectively. Weak induction (less than 0.5 log log10FC) of IFN-α and -β and ISG15 was observed in bronchial HAE at 37°C, except for the IFN-β induction by the recent subvariants XBB.1.5, EG.5.1 and JN.1.1 (0.65, 0.48 and 0.84 log10 FC, respectively).

**Figure 6.**
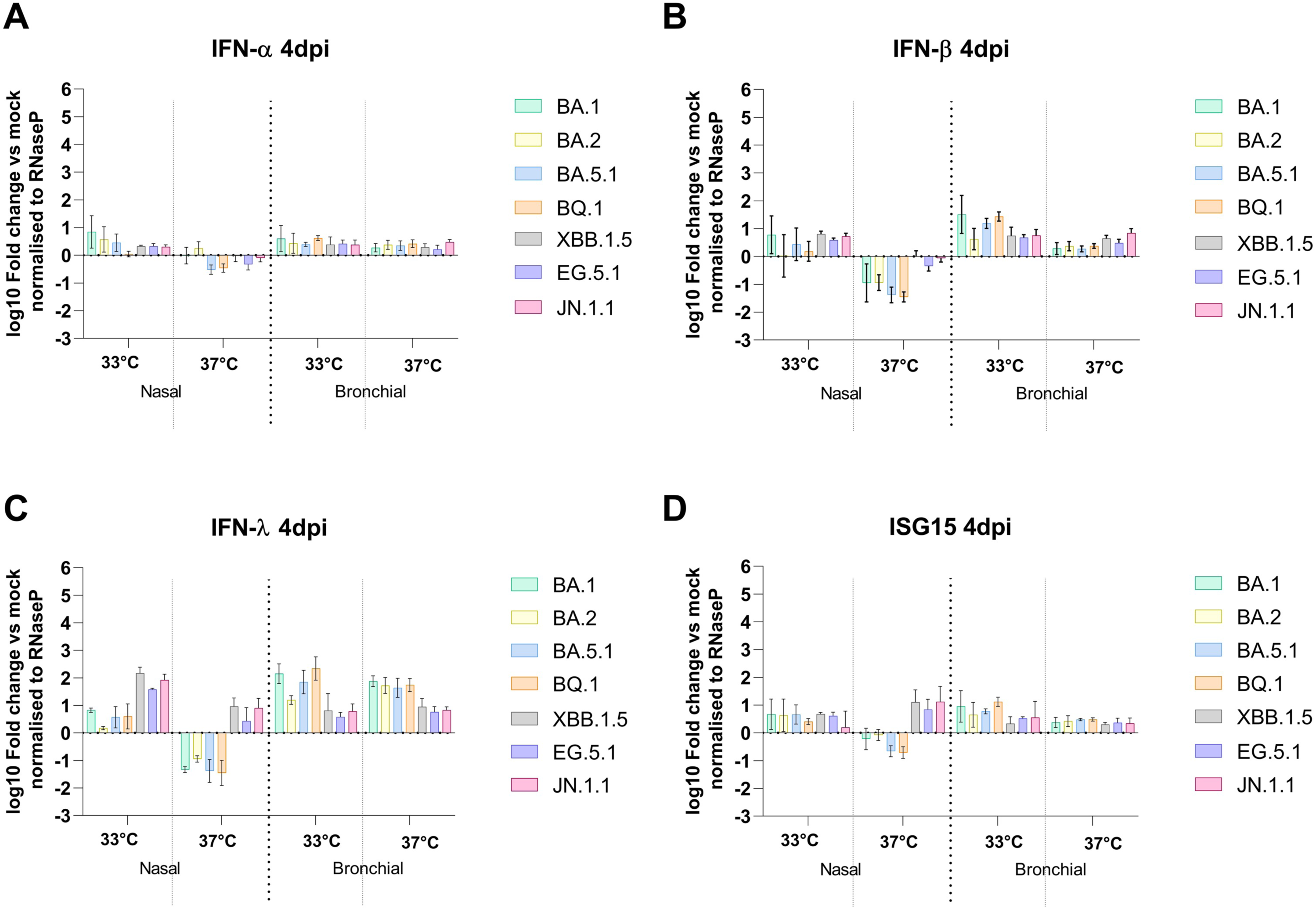
Induction of HAE intrinsic host response during by SARS-CoV-2 Omicron lineages. Nasal and Bronchial HAE infected with SARS-CoV-2 Omicron lineages BA.1, BA.2, BA.5.1, BQ.1, XBB.1.5, EG.5.1 and JN.1.1 at 33°C (the same shown Figure 1) were lysed at 96hpi. Induction of IFN-α (A) IFN-β (B), IFN-λ (C) and ISG15 (D) was assessed by semi-quantitative real time RT-PCR using intracellular RNA and expressed in fold change relative to non-infected and normalized to RNAseP. Data are represented as mean and SEM (n=3 donors), as for each cell origin.

IFN-λ showed the highest level of induction with all variants (except BA.2 in nasal HAE at 33°C) and the most pronounced variability between subvariants. While low induction was observed in BA.1-, BA.2- and BQ.1-infected nasal tissues at 33°C, XBB.1.5, EG.5.1 and JN.1.1 induced higher induction levels (2.17, 1.57- and 1.92-log10 FC, respectively). Inversely, the latter showed lower induction levels in bronchial HAE at 33°C (from 0.58 to 0.81 log10 FC versus from 1.19 to 2.34 log10 FC for BA.1, BA.2, BA.5.1 and BQ.1) and at 37°C from 0.76 to 0.95 log_10_ FC versus from 1.63 to 1.84 log_10_ FC for BA.1, BA.2, BA.5.1 and BQ.1). Induction of the main entry host factors, ACE-2 and TMPRSS2, was not enhanced in all conditions of Omicron lineages’ infections (**Figure S2**).

Altogether, our data showed, regardless of inter-donor variability, comparable replicative capacity in both tissue origins under both temperature conditions, although lower for BA.2 and slightly better for the most recent subvariants in nasal HAE.

Omicron lineages showed low induction of host responses, with an overall only slight difference between variants.

## Discussion

With the emergence of Omicron in late 2021, the COVID-19 pandemic has entered a new phase. It has previously been shown that Omicron, compared to earlier VOCs, could overcome immunity from various exposures, including prior infections and/or vaccinations, and showed a distinct phenotype in *ex-vivo* infections compared to previous variants [2]. In the continuous emergence of new Omicron lineages, intrinsic transmissibility and immune pressure are considered as the main drivers of viral evolution [24]. A range of studies have shown increasing escape from prior immunity for Omicron lineages, particularly those that are currently designated as VOIs such as XBB.1.5 like and BA.2.86.

Omicron BA.2.86, with its highly mutated spike carrying over 30 mutations compared to other Omicron lineages and a genetic distance that is comparable to that of the first Omicron lineages BA.1 to that of Delta, was initially suspected to have potentially enhanced immune escape properties [25]. Multiple neutralization studies showed similar or slightly diminished immunity evasion compared to other Omicron variants [10, 14, 26–33]. Despite its first detection in mid-2023, the prevalence of BA.2.86 remained low, possibly due to presumed lower viral fitness observed in cell culture studies and reduced pathogenicity in animal models compared to other Omicron lineages [14, 27, 34].

In line with clinical observations, previous studies including ours recently confirmed the faster but shorter replication of Omicron BA.1 compared to previous SARS-CoV-2 variants in nasal HAE [16]. We here confirmed this typical replication for BA.1 and extended this observation to more recent subvariants relative to Omicron, despite modest differences (especially with BA.2) in nasal and bronchial HAE models. A sustained phenotype of Omicron lineages has also been shown when looking at their intrinsic host response mainly involving IFN-λ. Even with an overall equal replicative capacity at upper and lower respiratory tract temperatures, as previously shown for Omicron but not the previous ancestral (B.1) and Alpha variants [35], little differences in IFN induction were observed at 37°C compared to 33°C. Higher replication efficiency in *in vitro* (Calu3) and *in vivo* (Balb/c mice) of BA.5, compared to BA.1, in lung tissues/cells have been reported [36].

More *ex-vivo* (explant) and *in vivo* (mice) studies suggested an association between the milder severity of Omicron lineages and their enhanced replication efficacy in the upper, compared to the lower, respiratory tract, in contrast to Delta variant [37–39]. One study found increased replicative capacity and infectivity of BA.5 in comparison to BA.1 and the ancestral virus in human nasal and airway organoids at 37°C [40]. In HAE, Zaderer *et al.* showed that, compared to the Deta variant, there was a decreased replication efficiency with Omicron subvariants BA.1, BA.2, BA.5 and BQ.1.1, and a superficial localization into the pseudostratified tissue with less pronounced anti-viral response [41]. The more recently emerging lineages, like the XBB, EG.5 and JN.1 benefit from an additional fitness advantage, as experimental data obtained with live virus in cell culture/primary cell cultures infection studies. We showed a small replication advantage and slightly higher IFN-λ response [19, 20] in nasal epithelia with these variants, in line with Planas *et al* [14].

In comparison with the parent lineage BA.2.86, JN.1 has an additional L455S substitution in the spike protein that was described to be associated with increased escape from humoral immunity as well as transmissibility [5, 11–15]. However, in contrast to its parental lineage BA.2.86, JN.1 rapidly outcompeted earlier variants in late 2023 and became domiant, associated with a wave of infections worldwide [11, 15]. Limited data on the immune escape assessment of JN.1 are available, but it demonstrated lower geometric mean neutralization titers and lower fold change values compared to earlier Omicron variants [15, 42].

The present study aimed at the assessment of emerging SARS-CoV-2 variants in an authentic virus neutralization assay against a panel of human sera and plasma. To add to the complexity of the underlying immunity, we used cohorts with immunity at both ends of the exposure spectrum, which reflects the complex situation in the population in the fifth year of the pandemic. Here we show that recently emerged SARS-CoV-2 Omicron variants, namely BQ.1, XBB.1.5, EG.5.1 and JN.1.1, display pronounced immune evasion to earlier Omicron variants, but that JN.1.1, does not show additional immune escape compared to EG.5.1. We also showed that earlier findings on hybrid immunity remain valid; specifically, hybrid immunity continues to offer higher neutralization titers compared to monovalent vaccination or natural infection. The use of authentic live virus isolates and FRNT_90_ to assess the neutralization titers of a large number of serum/plasma samples with a heterogeneous immunological background of the population adds strength to our findings. Indeed, according to the reported data, results may be different using a pseudovirus instead of a live virus, underscoring the importance of having data with authentic live viruses. It should be noted that our study has some limitations. First, we had a low number of sera that were available for the individual groups, especially for those that were exposed to more than one variant. It is important to note that vaccine sera from bivalent vaccines or updated vaccine formulations were not included in the analysis, which could impact the comprehensiveness of the findings regarding vaccine efficacy against emerging variants.

In summary, our data show that continuous assessment of newly evolving SARS-CoV-2, taking the different groups within the population into account, remains crucial to understand viral strategies to overcome existing immunity. In the case of JN.1, that showed a rapid global increase but no enhanced immune escape, other factors than immune escape seem to be the driving force behind this variant success. Collectively, this comparative study of the fitness and the immune escape capacity of the most relevant SARS-CoV-2 Omicron lineages, using pertinent cell models, authentic viruses and human specimens from immunized individuals, highlights the role of both virus fitness and adaptive immune response pressure on the evolution of Omicron lineages. It hence contributes to the better understanding of SARS-CoV-2 dynamics including its main driving forces as well as its phenotypical impact on viral properties.

## Acknowledgments

The authors acknowledge NIBSC for providing the VERO-E6-derived cells, Samuel Cordey for sequence analysis and Pascal Cherpillod for facilitating the work in the BSL-3 laboratory at HUG, staff of the laboratory of virology at the HUG for support, clinicians and technicians responsible for clinical cohorts, the population epidemiology of the HUG Unit for their help with participants’ recruitment and sample collection and all the donors.

## Authorship contributions

M.Bek., and M.E.L performed the experiments and conducted the analysis; K.A., C.A., P.S.R, D.D. and M.Bel helped to carry out some experiments; M.Bek, K.H.F, C.S.E., M.E.Z, S.S., I.G., O.P., M.Bel., and L.K. conducted the clinical studies and/or helped with clinical sample collection; I.E., M.Bek designed and supervised the study; M.Bek., M.E.L and I.E wrote the main manuscript. All authors have contributed to the final version of the manuscript.

## Figures

**Figure S1.**
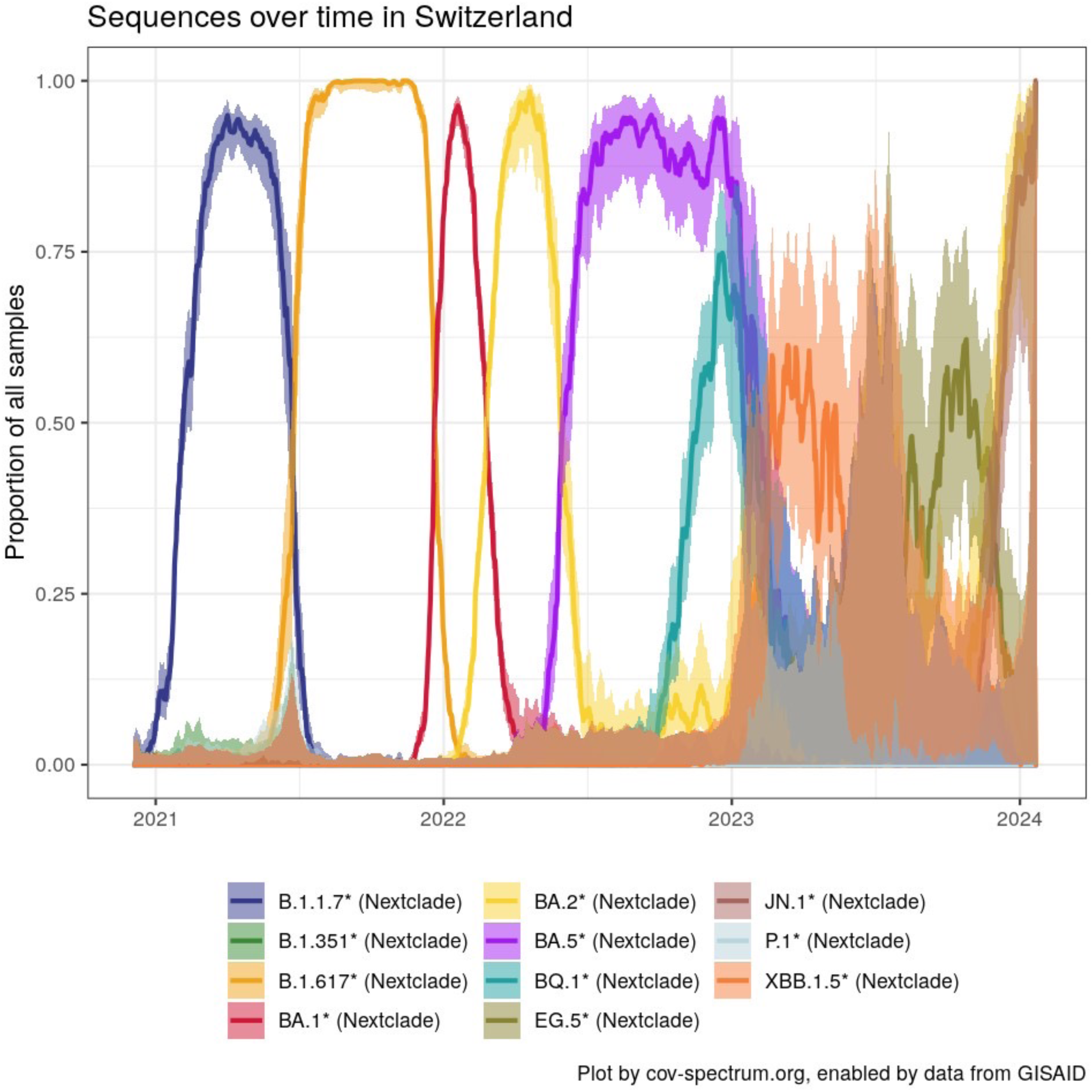
Trend of SARS-CoV-2 variants in Switzerland over time. From December 2020 until September 2023: from left to right, Alpha (B.1.1.7), Beta (B.1.351), Gamma (P.1), Delta (B.1.617), BA.1, BA.2, BA.5, BQ.1, XBB.1.5, EG.5.1 and JN.1.1 SARS-CoV-2 variants (https://cov-spectrum.org/explore/Switzerland/AllSamples/Past6M).

**Figure S2.**
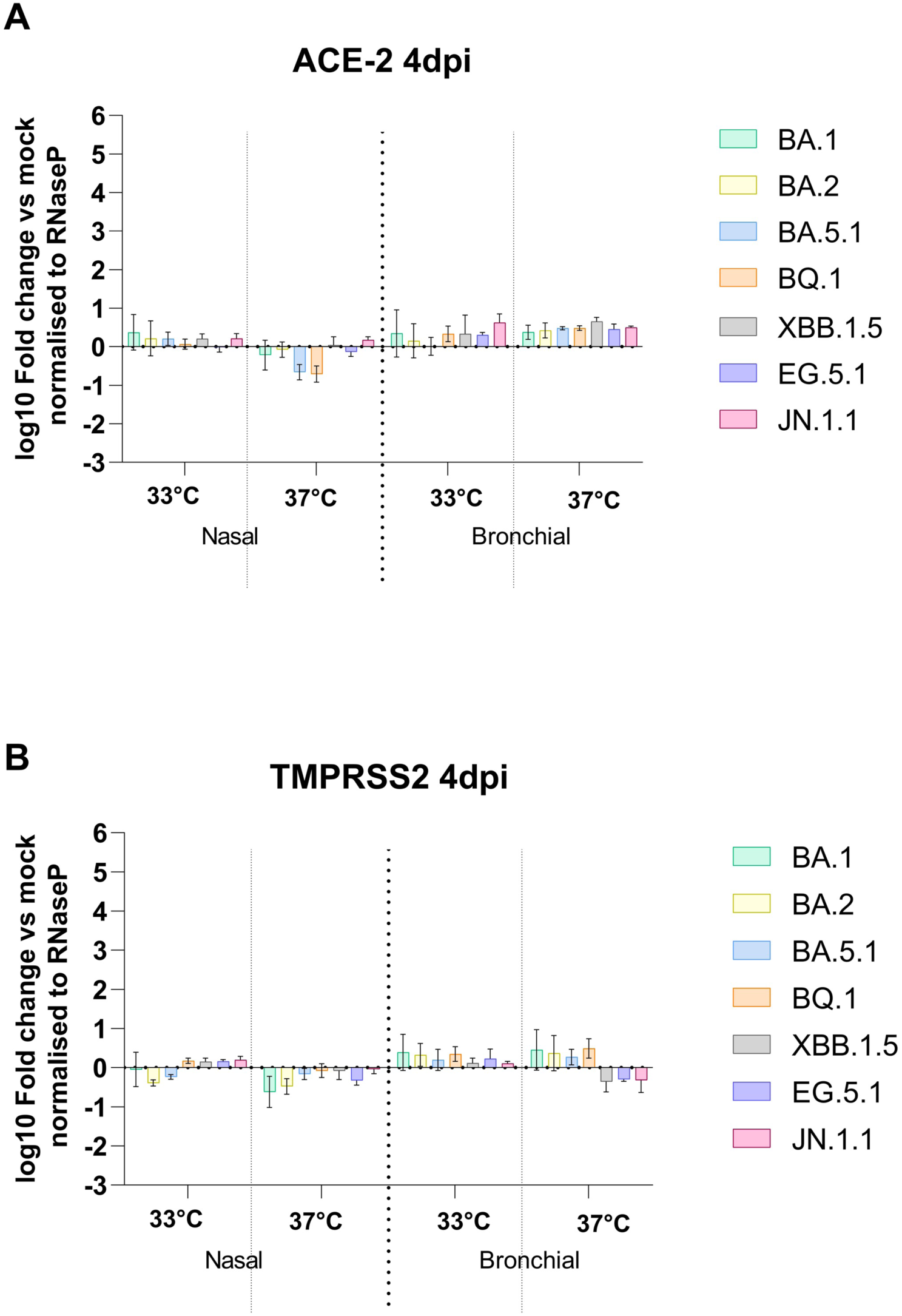
Induction entry host factor in HAE during infection by SARS-CoV-2 Omicron lineages. Nasal and Bronchial HAE infected with SARS-CoV-2 Omicron lineages BA.1, BA.2, BA.5.1, BQ.1, XBB.1.5, EG.5.1 and JN.1.1. at 33°C (as in Figures 5 and **6**) were lysed at 96hpi. Induction of host ACE-2 receptor (A) and TMPRSS2 protease (B), involved in SARS-CoV-2 entry during infection, was assessed by semi quantitative real time PCR using total RNA from the cell and expressed in fold change relative to non-infected and normalized to RNAseP. Data are represented as mean and SEM (n=3 epithelia from 3 different donors tested for each cell origin, nasal/bronchial).

